# Dissociation of Diffusion and Perfusion Responses After Intracerebral Hemorrhage

**DOI:** 10.64898/2026.01.09.698716

**Authors:** Xiuli Yang, Yuguo Li, Adnan Bibic, Zhiliang Wei

## Abstract

**Background and Purpose:** Intracerebral hemorrhage (ICH) triggers complex secondary injury processes that extend beyond hematoma formation. While structural and diffusion MRI are widely used to characterize tissue injury, the spatiotemporal evolution of cerebral perfusion after ICH, particularly in small-animal models, remains poorly defined. Here, we performed a longitudinal multiparametric MRI study to delineate the relationship between microstructural injury and cerebral perfusion following experimental ICH.

**Methods:** A collagenase-induced mouse model of ICH was studied longitudinally from baseline to 21 days post-stroke. Hematoma volume, tissue microstructure, and cerebral blood perfusion (CBP) were quantified using T2*-weighted MRI, diffusion-weighted imaging, and pseudo-continuous arterial spin labeling (pCASL) MRI, respectively. Apparent diffusion coefficient (ADC) and CBP were quantified in multiple brain regions from both ipsilateral and contralateral hemispheres and analyzed using linear mixed-effects models.

**Results:** Hematoma volume peaked acutely and gradually attenuated over time. ADC exhibited an early reduction largely confined to the striatum, followed by progressive recovery, consistent with localized cytotoxic edema and subsequent attenuation. In contrast, CBP showed a marked bilateral hypoperfusion during the acute phase, followed by a delayed perfusion increase that was spatially restricted to the ipsilateral striatum. Notably, significant contralateral perfusion alterations were observed despite minimal contralateral diffusion changes, indicating a dissociation between microstructural injury and vascular regulation.

**Conclusions:** Microstructural and perfusion responses after ICH follow distinct spatiotemporal trajectories. Whereas diffusion abnormalities are largely localized to the hemorrhagic core, perfusion disturbances extend bilaterally beyond the lesion site. These findings challenge the common assumption of contralateral physiological stability after focal hemorrhage and highlight the value of quantitative perfusion MRI for capturing systemic cerebrovascular responses that are not reflected by diffusion or anatomical measures alone.

## 1. Introduction

Intracerebral hemorrhage (ICH) represents a subtype of stroke caused by the rupture of cerebral blood vessels with extravasation of blood into the brain parenchyma or ventricular system [1]. ICH is characterized by a distinct pathological cascade in comparison with the ischemic stroke, including immediate mechanical tissue disruption, exposure of neural tissue to blood-derived toxic products, and rapid neurological deterioration [1, 2]. Despite of its relatively smaller proportion among all stroke cases (approximately 10-20%) [3], ICH conveys disproportionately high mortality, morbidity, and long-term disability [4]. Survivors often suffer from persistent neurological and cognitive impairments, which cannot be effectively alleviated after post-stroke rehabilitation and reflects complex secondary injuries, such as inflammation, edema, blood-brain barrier disruption, and vascular dysfunction [5]. Unfortunately, effective therapeutic options for ICH remain limited in clinic, underscoring the need to better understand underlying pathophysiology and to develop quantitative biomarkers capable of tracking injury evolution and informing therapeutic intervention.

Clinical ICH research has focused on phenotype characterization, risk factor identification (such as hypertension and amyloid angiopathy), and acute management strategies such as blood pressure control, surgical hematoma evacuation, and hemostatic therapies [6]. Large clinical trials have evaluated interventions to limit hematoma expansion and improve functional outcomes [7], yet progress toward effective, evidence-based neuroprotective treatments has been modest. Parallel preclinical work focused on dissecting mechanisms of secondary injury, including edema formation, excitotoxicity, inflammatory cascades, blood-brain barrier disruption, and molecular/cellular contributors to tissue damage [8–10]. Despite of these clinical and preclinical advancements, a critical gap remains in linking tissue-level pathophysiology with noninvasive biomarkers that can monitor pathological progression and therapeutic response in ICH. In recent years, there is a growing interest in leveraging advanced imaging modalities such as quantitative MRI to characterize cerebral blood flow, metabolism, vascular reactivity, and blood-brain barrier integrity in cerebrovascular diseases [11–15], in the expectation of unravelling biological underpinnings of neurofunctional impairment, identifying novel targets for intervention and long-term tracking, and moving descriptive characterization toward predictive insights.

Previous studies have characterized hematoma expansion and microstructural alterations after ICH using structural and diffusion MRI [16, 17], providing important insights into tissue injury and edema evolution. However, far less is known about the longitudinal changes in cerebral perfusion after ICH, particularly in mouse models, where noninvasive and quantitative assessments remain technically challenging. In experimental and clinical neuroimaging studies of ICH, the contralateral hemisphere is commonly assumed to remain physiologically stable and is therefore frequently used as an internal reference for normalization or control comparisons [10, 16, 18]. However, whether this assumption holds for cerebrovascular regulation following ICH has not been systematically examined. Given the prominent roles of systemic inflammation [19] and autonomic dysregulation [20] after hemorrhage, contralateral perfusion may itself be dynamically altered, potentially confounding interpretation of ipsilateral injury metrics.

In this study, we performed a multiparametric MRI characterization of a collagenase-induced mouse ICH model [21] from baseline to 21 days post-stroke. While hematoma volume and apparent diffusion coefficient (ADC) measurements were used to contextualize tissue injury, we leveraged pseudo-continuous arterial spin labeling (pCASL) MRI to quantify cerebral blood perfusion (CBP) longitudinally and to delineate hemisphere-specific perfusion dynamics following hemorrhagic injury. By integrating established structural and microstructural markers with perfusion imaging, this work aims to provide a physiological perspective on post-ICH evolution that extends beyond conventional anatomical assessments.

## 2. Methods

### 2.1 General

The experimental protocols for this study were approved by the Johns Hopkins Medical Institution Animal Care and Use Committee and conducted in accordance with the National Institutes of Health guidelines for the care and use of laboratory animals. Data reporting complied with the ARRIVE 2.0 guidelines. All procedures were carefully designed to minimize discomfort and stress to the animals. Fourteen male C57BL/6 mice were included, with ages of 3 months (*n* = 4), 6 months (*n* = 3), 16 months (*n* = 3), and 22 months (*n* = 4). Multiple age cohorts were included to better reflect the age heterogeneity observed in clinical populations and enhance the generalizability of results. Mice were housed in a quiet environment under a 12-h light/dark cycle with ad libitum access to food and water.

Intracerebral hemorrhage (ICH) was induced using previously established protocols [21]. Briefly, mice were anesthetized with isoflurane (70% N_2_ and 30% O_2_; 4% induction, 2% maintenance) and secured in a stereotaxic frame. Following a 1 cm sagittal scalp incision, collagenase VII-S (0.075 U in 0.5 μL saline; Sigma, St. Louis, MO, USA) was infused into the right striatum (AP: +0.6 mm, ML: +2.1 mm, DV: −3.0 mm relative to bregma) at a rate of 0.1 μL/min using a micro-infusion pump (Hamilton, Reno, NV, USA) for 5 mins, the needle was left in place for another 5 mins and then removed. Body temperature was regulated at 37.0 ± 0.5°C. The burr hole was sealed with bone wax, and the skin incision was closed. To ensure model consistency, mice experiencing mortality within 24 hours were excluded from the study.

### 2.2 MRI experiments

All MRI experiments were performed on an 11.7T Bruker Biospec system (Bruker, Ettlingen, Germany) with a horizontal bore and an actively shielded pulse-field gradient (maximum intensity of 0.74 T/m). Images were acquired using a 72-mm quadrature volume resonator as the transmitter, and a four-element (2×2) phased-array coil as the receiver. Magnetic field homogeneity over the mouse brain was optimized using global shimming (up to the second order) based on a pre-acquired subject-specific field map. Inhalational isoflurane, delivered with medical air (21% O_2_ and 78% N_2_) or hypercapnic gas (5% CO_2_, 21% O_2_, and 74% N_2_) at a flow rate of 0.75 L/min, was used to minimize stress and motion in the mice. Anesthesia induction was achieved with 1.5-2.0% isoflurane for 15 minutes. At the 10^th^ minute under induction, the mouse was placed onto a water-heated animal bed with temperature control and positioned with a bite bar, a pair of ear pins, and a custom-built 3D-printed holder before entering the magnet [22]. After induction, the isoflurane concentration was reduced to 1.0% for maintenance during the MRI scans. Respiration was observed during the experiments using an MRI-compatible monitoring system (SA Instruments, Stony Brook, NY, USA). Isoflurane dose was adjusted as needed to maintain a respiration rate of 70–120 breaths per minute [23]. Mice were scanned in a randomized order, following a previously reported scheme [24]. A subset of hemorrhagic mice was examined longitudinally at multiple time points: pre-stroke (Day 0), and 1, 3, 7, 14, and 21 days post-stroke (Days 1, 3, 7, 14, and 21, respectively; Figure 1). The final sample sizes at these time points were *n* = 6, 8, 11, 4, 4, and 4 (37 scan sessions totally), respectively. Multiparametric MRI characterizations, including gradient echo (GRE) MRI, diffusion-weighted imaging (DWI), and pCASL MRI, were performed (Figure 1) to examine the longitudinal changes in hematoma volume, ADC, and CBP.

**Figure 1.**
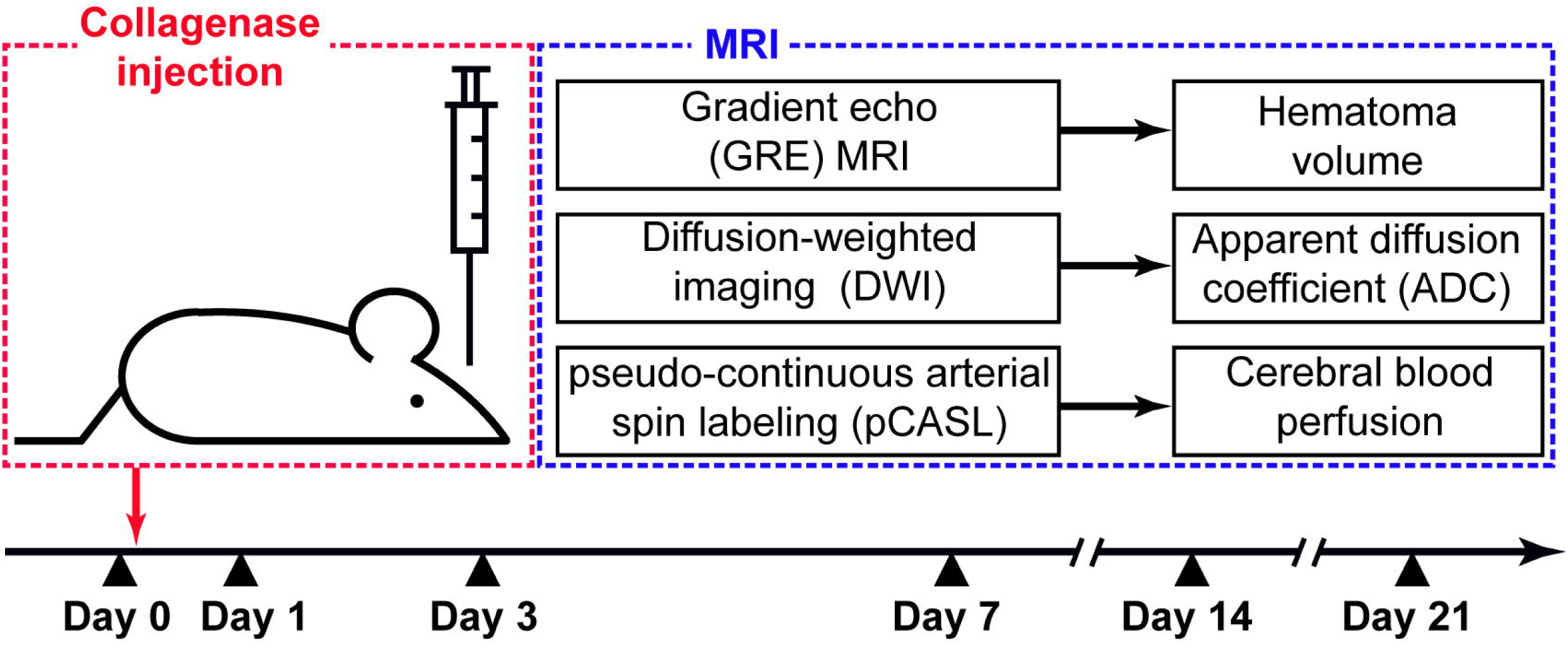
Experimental design and MRI protocol. The intracerebral hemorrhage (ICH) model was induced by intra-striatal collagenase injection. A baseline MRI scan (Day 0) was acquired for reference, followed by post-ICH imaging at five time points (Days 1, 3, 7, 14, and 21). The MRI protocol included: T_2_*-weighted gradient-echo (GRE) imaging for hematoma volume assessment; diffusion-weighted imaging (DWI) for microstructural characterization indexed by the apparent diffusion coefficient (ADC); and pseudo-continuous arterial spin labeling (pCASL) for quantification of cerebral blood perfusion (CBP).

Key imaging parameters of GRE MRI were [25]: repetition time (TR) = 200 ms; echo time (TE) = 12.0 ms; field of view = 15 (left–right) × 7.5 (ventral–dorsal) × 4.8 (rostral–caudal) mm^3^; matrix size = 256 × 128 × 24; spatial resolution = 59 × 59 × 200 µm^3^; receiver bandwidth = 110 kHz; partial-Fourier factor = 0.75 along the phase-encoding direction; and scan duration = 7.7 min with three-dimensional acquisition.

Key imaging parameters of DWI scan were as follows [23]: TR/TE = 1000/17.0 ms; field of view = 15.0 × 7.5 × 4.8 mm^3^; matrix size = 128 × 64 × 24; spatial resolution = 117 × 117 × 200 µm^3^; encoding direction = 6; *b* = 650 s/mm2; three repetitions of non-diffusion-weighted (*b* = 0 s/mm^2^) image; receiver bandwidth = 300 kHz; partial Fourier factor = 0.63 along the phase-encoding direction; segment number of spin-echo echo-planar imaging (EPI) = 2; and scan duration = 7.2 min with three-dimensional acquisition.

A two-scan pCASL scheme developed to improve the robustness of labeling against magnetic-field inhomogeneity was utilized in this study [26]. A pre-scan was first performed to optimize the phase offsets of labeling pulses for both control and label scans [27]. Subsequently, pCASL scans focusing on perfusion imaging were performed under the following parameters [28]: repetition time (TR) = 3000 ms; echo time (TE) = 11.8 ms; field of view = 15 × 15 mm^2^; slice thickness = 0.75 mm; interslice gap = 0.25 mm; matrix size = 96 × 96; number of slices = 12; receiver bandwidth = 300 kHz; labeling duration = 1800 ms; inter-labeling pulse delay = 1.0 ms; labeling pulse width = 0.4 ms; labeling plane thickness = 1.0 mm; labeling pulse flip angle = 40°; mean gradient = 0.01 T/m (label scans) and 0 T/m (control scans); post-labeling delay (PLD) = 300 ms; signal average = 25; and scan duration = 5.0 min using a two-segment spin-echo EPI acquisition. Labeling efficiency was measured using the bidirectional crusher gradient method [27, 28].

### 2.3 Data processing

All data processing was performed using custom MATLAB scripts (MathWorks, Natick, MA). GRE images were analyzed in a slice-by-slice manner to define hematoma boundaries with reference to normal anatomy from the Allen Brain Atlas. The total hematoma volume was estimated by summing voxel counts across slices and multiplying by the voxel volume.

Ipsilateral and contralateral regions of interest (ROIs) were manually delineated on the non-diffusion-weighted images of DWI to encompass the midbrain, isocortex, hippocampus, thalamus, hypothalamus, and striatum. Mean ADC values were calculated to characterize region-specific microstructural properties.

Processing of the pCASL data followed established procedures [23]. CBP was calculated as 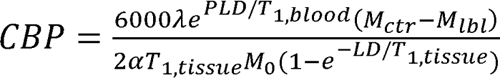 [26], where A denotes the brain-blood partition coefficient (0.89 mL/g [29]); T_1,blood_ is the longitudinal relaxation time of blood (2813 ms at 11.7 T [30]); M_ctr_ and M_lbl_ are the control and label images; a is the labeling efficiency; M_0_ denotes the equilibrium magnetization (obtained by scaling the control image [26]); T_1,tissue_ is the longitudinal relaxation time of tissue (1900 ms [31]); and LDdenotes labeling duration. ROI analyses, analogous to those used for DWI processing, were performed to assess regional CBP patterns.

### 2.4 Statistical analyses

Linear mixed-effects (LME) models were used to examine the dependence of hematoma volume, ADC, and CBP on time points. Day_ICH_ denoted the time points of ICH and was treated as a continuous predictor; Day^2^ was included to explore the nonlinear pattern when applicable; Side was a categorical predictor denoting the ipsilateral and contralateral hemispheres of ICH; age was included as a co-variate; and a random intercept (E_mouse_) was included for mouse. LME models were fit by restricted maximum likelihood (REML), with degrees of freedom estimated using the residual method. Effect estimate (denoted as β), 95% confidence interval (CI), and *p*-value were reported. Details of all statistical analyses were summarized in Supporting Table S1. Measurement values were presented as mean ± standard deviation. Bonferroni correction was applied to control for multiple comparisons, and significance was defined as *p* < 0.05 divided by the number of tests. For example, a significance threshold of *p* < 0.008 was applied to analyses across the six brain regions. In addition, *p*-values were presented with scientific notation when appropriate, for example, 1.950E-07 represented 1.950 × 10^-7^.

## 3. Results

### 3.1 Time-dependent attenuation of hematoma following ICH

Regarding ICH, the hematoma refers to the localized intraparenchymal accumulation of extravasated blood resulting from vascular rupture. On GRE MRI, it can be readily identified as a hypointense region arising from susceptibility effects of blood products within the brain parenchyma, as illustrated in a representative mouse (Figure 2a). Hematoma volume at baseline (i.e., Day 0) was zero (Figure 2b). Including Day 0 in analyses would disproportionately affect model fitting and mask temporal trends of the post-stroke phase, which represented the formation and attenuation of hematoma (Figure 2b). Therefore, Day 0 was excluded in the analyses of hematoma volume using LME models.

**Figure 2.**
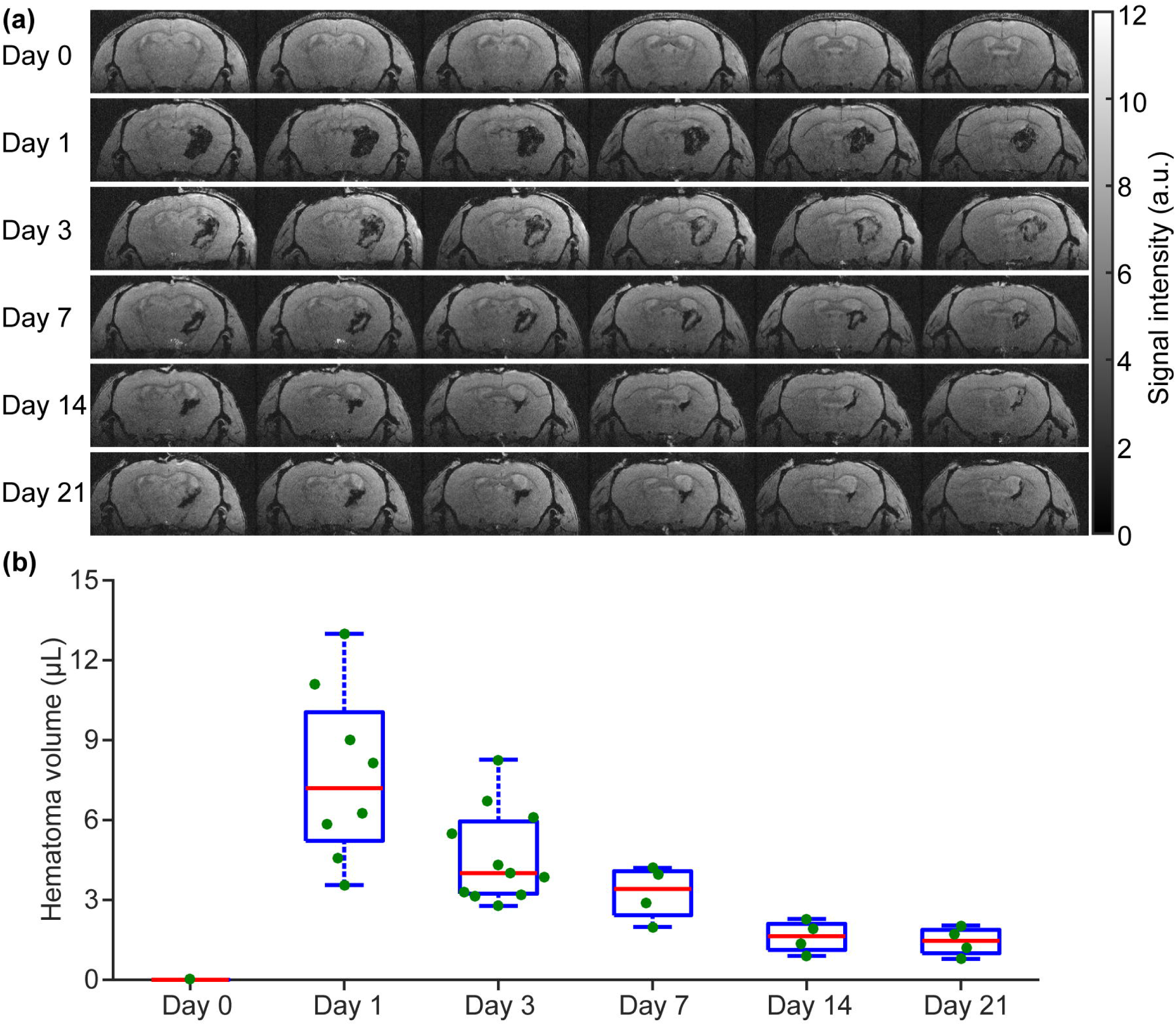
Temporal evolution of hematoma volume following ICH. (a) Representative longitudinal T_2_*-weighted images acquired from a single mouse across the indicated time points. (b) Hematoma volume plotted as a function of time. In the boxplots, the central red line denotes the median; the upper and lower edges of the box represent the 75^th^ and 25^th^ percentiles, respectively; and the whiskers extend to the minimum and maximum values excluding outliers.

Hematoma volume (denoted as V_hema_) exhibited a significant dependence on post-ICH time (β = −0.24 µL/day, *p* = 1.950E-07) with peak volume observed at Day 1 (Figure 2b). A subsequent analysis confirmed a nonlinear relationship between hematoma volume and post-ICH time, as evidenced by a significant quadratic term (β = 0.02 µL/day^2^, *p* = 0.002). The mean hematoma volumes were 7.68 ± 3.25, 4.66 ± 1.76, 3.25 ± 1.03, 1.61 ± 0.61, and 1.44 ± 0.55 mm^3^ at 1, 3, 7, 14, and 21 days post-ICH, respectively, indicating an 81.3% reduction in hematoma volume by Day 21 relative to the peak at Day 1.

### 3.2 Microstructural tissue changes after ICH revealed by ADC

Representative ADC maps illustrate temporal changes in microstructural diffusivity across multiple brain regions (Figure 3a), highlighting an initial ADC hypointensity after ICH induction and a subsequent progressive increase over time (indicated by red arrows). Motivated by the hematoma volume analysis, data were categorized into two temporal phases, i.e., acute (Day 0 and Day 1) and post-acute (Days 1, 3, 7, 14, and 21), and analyzed accordingly.

**Figure 3.**
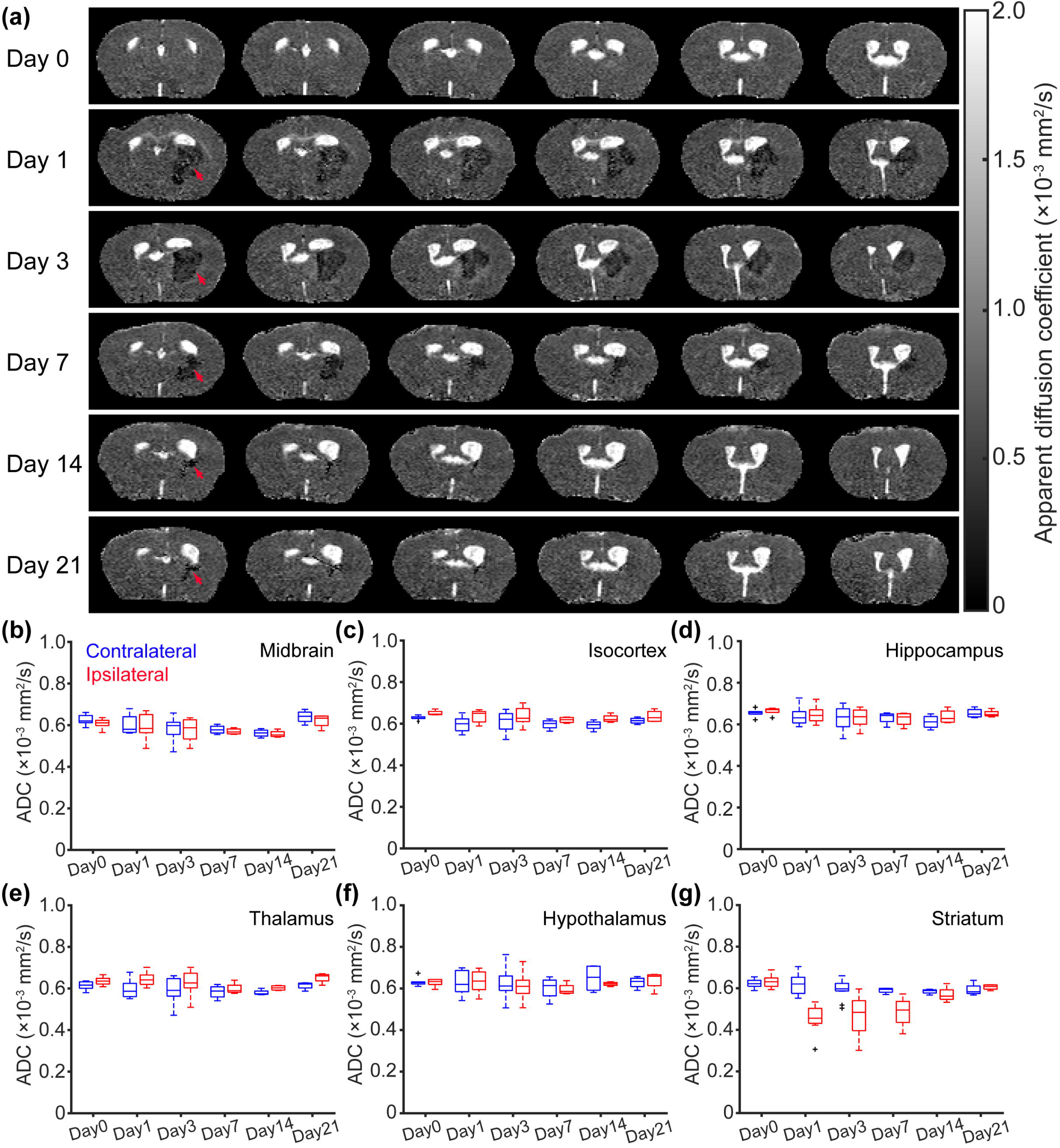
Temporal evolution of tissue diffusivity following ICH. (a) Representative longitudinal apparent diffusion coefficient (ADC) maps acquired from a single mouse across the indicated time points. (b–g) ADC plotted as a function of time in the midbrain, isocortex, hippocampus, thalamus, hypothalamus, and striatum, respectively. Contralateral and ipsilateral hemispheric data are shown in blue and red, respectively. In the boxplots, crosses denote outliers.

To examine whether ICH induced contralateral changes in ADC, LME analyses were performed using contralateral data only. At the acute stage, a significant Day_ICH_ effect was observed in the contralateral isocortex (β = −40.88 × 10^-6^ mm^2^/s/day, *p* = 0.002), whereas no significant effects were detected in other contralateral regions (*p* ≥ 0.013). These findings suggest that ICH may induce diffusivity alterations in the contralateral hemisphere. Potential mechanisms underlying the observed ADC decrease in the contralateral isocortex would be discussed later. At the post-acute stage, post-ICH time did not significantly affect contralateral ADC in any of the examined regions (*p* ≥ 0.160).

At the acute phase, the ipsilateral hemisphere exhibited a distinct pattern compared with the contralateral side. Significant Day_ICH_ effects were observed in both the ipsilateral thalamus (β = −5.41× 10^-6^ mm^2^/s/day, *p* = 0.003) and striatum (β = −183.41 × 10^-6^ mm^2^/s/day, *p* = 4.792E-05). The marked ADC reduction in the striatum is consistent with the development of cytotoxic edema, in which failure of energy-dependent ion pumps (primarily Na^+^/K^+^-ATPase) led to intracellular water accumulation. The motion of intracellular water molecules was subsequently restricted by cell membranes and intracellular organelles, resulting in reduced diffusivity. The thalamus is anatomically adjacent to the striatum, which served as the primary site of ICH induction. Therefore, the observed decrease in thalamic ADC is likely attributable to unintended extension of the hemorrhagic lesion beyond the targeted striatal region. At the post-acute phase, striatal ADC demonstrated a significant positive dependence on post-ICH time (β = 7.14 × 10^-6^ mm^2^/s/day, *p* = 3.384E-05), indicating progressive attenuation of cytotoxic edema.

We further examined the interaction between Side (contralateral vs. ipsilateral) and Day_ICH_. A significant interaction effect was detected exclusively in the striatum at both the acute (β = −177.75 × 10^-6^ mm^2^/s/day, p = 8.440E-05) and post-acute (β = 9.01 × 10^-6^ mm^2^/s/day, p = 1.198E-07) phases, indicating that the striatum exhibited the largest inter-hemispheric difference in the temporal trajectory of ADC.

Taken together, these findings indicate that the diffusivity cascade following ICH is largely confined to the induction region, namely the striatum, with minimal propagation to remote brain regions.

### 3.3 Temporal trajectory of brain perfusion following ICH

The time-dependent diminishing and recovery of brain perfusion was highlighted by the arrows in the CBP maps (Figure 4a).

**Figure 4.**
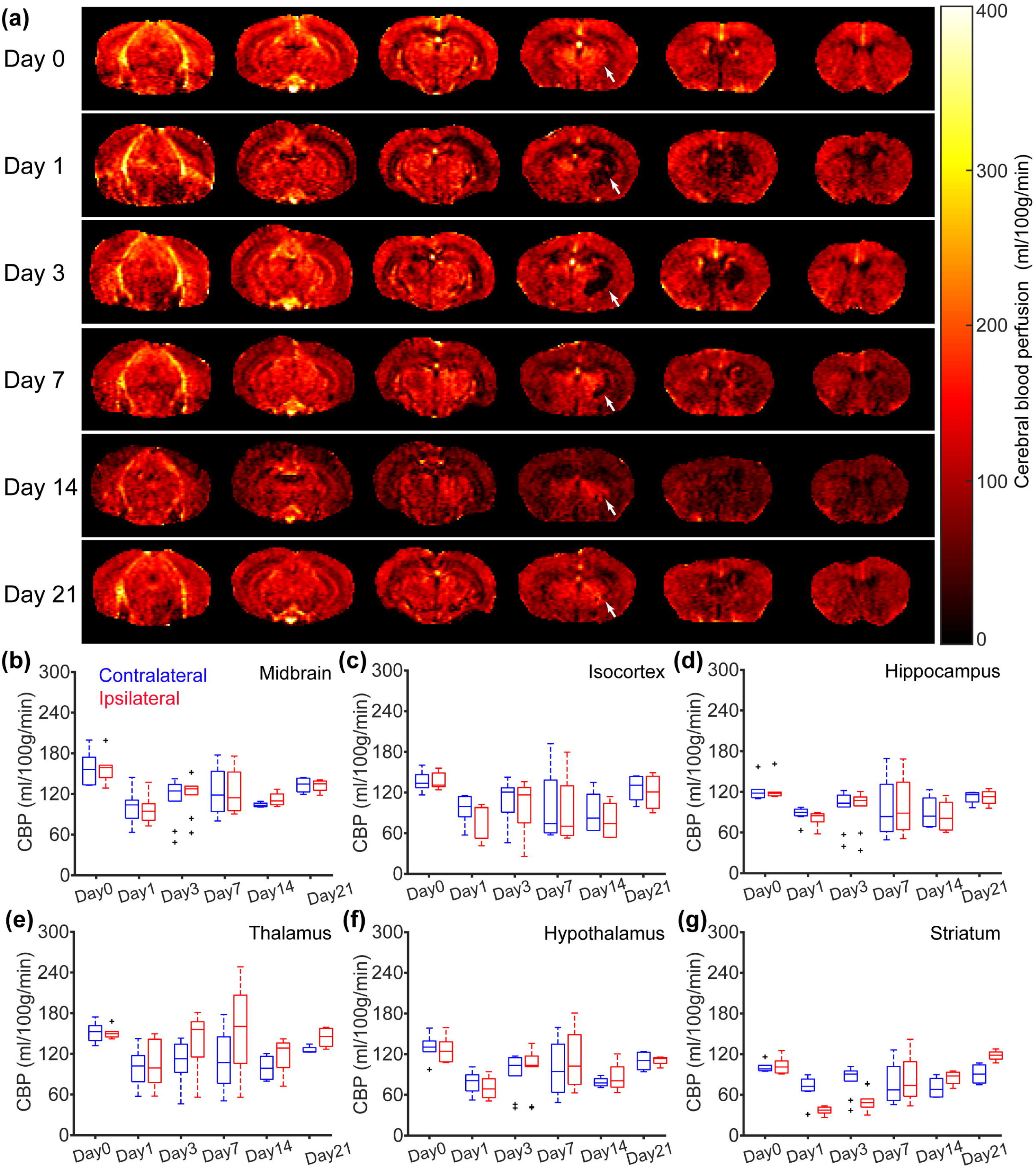
Temporal evolution of cerebral blood perfusion (CBP) following ICH. (a) Representative longitudinal CBP maps acquired from a single mouse across the indicated time points. (b–g) CBP plotted as a function of time in the midbrain, isocortex, hippocampus, thalamus, hypothalamus, and striatum, respectively. Contralateral and ipsilateral hemispheric data are shown in blue and red, respectively. In the boxplots, crosses denote outliers.

At the acute phase, significant Day_ICH_ effects were observed in contralateral CBP across multiple brain regions, including the midbrain (β = −57.93 ml/100g/min/day, *p* = 9.690E-04), isocortex (β = −38.95 ml/100g/min/day, *p* = 3.035E-04), hippocampus (β = −36.31 ml/100g/min/day, *p* = 9.573E-05), thalamus (β = −52.56 ml/100g/min/day, *p* = 0.001), hypothalamus (β = −50.38 ml/100g/min/day, *p* = 2.687E-05), and striatum (β = −30.82 ml/100g/min/day, *p* = 0.002). Consistent results were obtained from separate analyses of ipsilateral CBP, which demonstrated significant Day_ICH_ effects in the midbrain (β = −61.91 ml/100g/min/day, *p* = 1.766E-04), isocortex (β = −54.33 ml/100g/min/day, *p* = 2.344E-04), hippocampus (β = −44.00 ml/100g/min/day, *p* = 3.192E-05), thalamus (β = −46.97 ml/100g/min/day, *p* = 7.720E-03), hypothalamus (β = −55.24 ml/100g/min/day, *p* = 2.619E-05), and striatum (β = −66.92 ml/100g/min/day, *p* = 6.482E-08). Collectively, these findings indicate a widespread reduction in CBP during the acute phase following ICH induction. This global perfusion decrease is plausibly related to systemic inflammatory responses and cytokine-mediated vasoactive effects triggered by the hemorrhagic insult.

At the post-acute phase, a significant Day_ICH_ effect was observed exclusively in the ipsilateral striatal CBP (β = 3.75 ml/100g/min/day, *p* = 1.042E-08), whereas no significant effects were detected in other contralateral or ipsilateral regions (*p* ≥ 0.236). These findings indicate a gradual recovery of CBP localized to the ipsilateral striatum.

When examining the interaction effect between Side and Day_ICH_, significant interaction effects were identified exclusively in the striatum at both the acute (β = −35.04 ml/100g/min/day, *p* = 3.290E-04) and post-acute (β = 3.31 ml/100g/min/day, *p* = 5.527E-07) phases. These results indicate that the striatum exhibited the most pronounced inter-hemispheric difference in the temporal evolution of CBP among the examined brain regions.

CBP results indicated a widespread hypoperfusion stress following the ICH insult. In brain regions intrinsically associated with high neuronal activity, such as the isocortex, this perfusion reduction may increase vulnerability to hypometabolism and subsequent cytotoxic edema, consistent with the observed decrease in ADC in the contralateral isocortex.

Collectively, CBP alterations following ICH were not confined to the induction region but extended across a broad range of contralateral and ipsilateral brain regions.

## 4. Discussion

In this study, we performed a longitudinal characterization of microstructural integrity and cerebral perfusion in a collagenase-induced mouse model of ICH. A key finding is the dissociation between microstructural recovery and vascular regulation following ICH. Diffusion-derived changes, indexed by ADC, were largely confined to the hemorrhage induction site, whereas perfusion alterations, measured by CBP, extended beyond the lesion core to involve widespread bilateral brain regions. This dissociation underscores the importance of interpreting specific pathophysiological alterations within a broader physiological context and highlights the value of multiparametric MRI for disentangling overlapping yet non-identical injury mechanisms. ADC and CBP analyses converged on the striatum as the region exhibiting the most pronounced and persistent hemisphere-specific effects. This regional specificity likely reflects the direct vascular perturbation at the induction site. Beyond that, contralateral alterations were noticed in both ADC and CBP. These findings call into question the widespread practice of contralateral normalization assuming that the contralateral hemisphere remains physiologically stable after focal hemorrhagic injury. Such an assumption is the methodological basis for implementing normalization analyses or using contralateral values as controls. Systemic inflammatory signaling [19] and altered autonomic regulation [20] etc. can contribute to bilateral physiological perturbations following ICH. Consequently, normalization of ipsilateral measurements to contralateral values, while useful for emphasizing asymmetry, may obscure meaningful global or bilateral effects. By analyzing absolute values in both hemispheres, the present study provides a more complete depiction of post-ICH physiological dynamics.

While hematoma volume and diffusion metrics have been extensively used to characterize tissue injury after ICH [10, 16, 32], noninvasive assessment of cerebral perfusion in small-animal hemorrhagic models remains limited. The pCASL MRI is a widely used non-contrast technique for CBP mapping; however, accurate estimation of labeling efficiency in small animals is technically challenging due to the substantially smaller arterial geometry compared with humans. In this study, we leveraged a recently developed bidirectional crusher gradient method to improve the accuracy of CBP quantification. Our findings demonstrate the feasibility and utility of quantitative perfusion imaging for capturing dynamic CBP responses following ICH. By providing a physiological dimension that complements conventional anatomical and diffusion-based assessments, pCASL MRI may serve as a valuable tool for evaluating vascular-targeted interventions and recovery processes in hemorrhagic stroke.

The observed temporal trajectories of hematoma volume and ADC are consistent with a previous report examining 3, 7, and 28 days post-ICH [16], reflecting the biological processes of hematoma formation and subsequent attenuation. A previous rat study using laser Doppler imaging have reported a decrease in relative cerebral blood flow during the first 90 minutes following ICH induction [10]. CBP in the hemorrhagic core was found to decrease significantly relative to contralateral regions at 1, 7, and 14 days post-ICH [18]. Extending these observations, our study demonstrates that hypoperfusion is not restricted to the hemorrhagic site or cortical regions but is instead widespread and persists over a substantially longer time window, on the order of days after ICH.

A competing mechanism that can lead to ADC elevation is vasogenic edema, in which circulating neuroinflammatory cytokines disrupt the blood-brain barrier, permitting increased water extravasation into the perivascular and extracellular spaces [32]. The resulting expansion of the extracellular compartment facilitates greater water mobility, thereby increasing diffusivity (i.e., higher ADC). In the present study, the observed increase in striatal ADC during the post-acute phase is more likely attributable to attenuation of cytotoxic edema rather than the development of vasogenic edema, as comparable ADC increases were not detected in other brain regions. This regional specificity argues against a dominant role of systemically circulating cytokines, which would be expected to induce more widespread BBB perturbation.

CBP changes exhibited a biphasic temporal profile characterized by an initial hypoperfusion affecting both hemispheres, likely reflecting systemic and cerebrovascular responses to acute hemorrhage [33–36], including elevated intracranial pressure, sympathetic activation, inflammatory cytokine release, and transient impairment of vascular tone. Previous studies have shown that neuroinflammation, as indicated by sustained increases in Iba1^+^ microglia/macrophages and GFAP^+^ astrocytes, can persist for up to 28 days following ICH [16]. Concurrently, neuroinflammatory processes have been linked to cerebral hypoperfusion across multiple brain regions [37]. Together, these observations suggest that persistent neuroinflammation is likely a major contributor to the prolonged hypoperfusion observed after ICH. On the other hand, a perfusion increase emerged that was spatially restricted to the ipsilateral striatum, suggesting a localized vascular process rather than a global restoration of perfusion. This delayed perfusion elevation may reflect reactive hyperemia, impaired autoregulatory control, or remodeling of the perihematomal microvasculature [38–40].

The findings of this study should be interpreted with several limitations. First, only male mice were included to minimize potential confounding effects of estrogen; therefore, future studies incorporating female mice are warranted to examine possible sex-dependent differences. Second, the overall sample size was moderate. It should be noted that this study primarily focused on temporal trajectories rather than between-group comparisons. The longitudinal design, emphasizing within-subject temporal evolution, enhanced statistical power for detecting dynamic changes. Follow-up studies with larger cohorts will be necessary to further validate and generalize the present findings. Finally, the current work focused on multiparametric MRI characterization of hematoma volume, tissue diffusivity, and cerebral perfusion. Future studies incorporating complementary approaches such as metabolomics [41] or magnetic resonance spectroscopy [42] may provide additional molecular-level insights into the underlying pathophysiology.

## 5. Conclusion

Post-hemorrhage physiological evolution is characterized by distinct and regionally heterogeneous trajectories of tissue microstructure and cerebral perfusion. Whereas diffusion abnormalities are largely confined to the hemorrhagic core, perfusion disturbances extend bilaterally beyond the lesion site, challenging the assumption of contralateral physiological stability after focal intracerebral hemorrhage. Together, these findings underscore the value of multiparametric physiological imaging for capturing systemic cerebrovascular responses that are not reflected by lesion-centric metrics alone.

## CRediT Author Contributions Statement

X.Y.: Conceptualization; Methodology; Investigation; Validation; Writing – Original Draft Preparation. Y.L. & A.B.: Investigation; Writing – Review & Editing. Z.W.: Conceptualization; Methodology; Software; Resources; Data Curation; Funding Acquisition; Writing – Original Draft Preparation; Writing – Review & Editing.

## Acknowledgements

This work did not receive any external funding. The authors would like to thank Kazi Akhter for assistance during the data collections.

## Data Availability Statement

All data supporting the findings of this study can be obtained from the corresponding author upon reasonable request.

## Abbreviations

ADC: apparent diffusion coefficient;
CBP: cerebral blood perfusion;
CI: confidence interval;
DWI: diffusion weighted imaging;
GRE: gradient echo;
ICH: intracerebral hemorrhage;
LME: linear mixed-effects;
pCASL: pseudo-continuous arterial spin labeling;
PLD: post-labeling delay;
ROI: region of interest;
TE: echo time;
TR: repetition time;

