## Supplemental figure for "Dissociation of Diffusion and Perfusion Responses After Intracerebral Hemorrhage"

### Supplemental material

#### 1. Table S1 Summary of statistical analyses.

$\varepsilon_{mouse}$ denotes the random intercept included for mouse. MB: midbrain; ISO: isocortex; HPC: hippocampus; TH: thalamus; HY: hypothalamus; STR: striatum. Significant effects are shown in bold red.

| Statistical models | Effect estimate (β) | 95% Confidence interval | *p*-value[g] |
|---|---|---|---|
| *Section A: Hematoma volume ($V_{hema}$)* | | | |
| $V_{hema} \sim Day_{ICH} + age + \varepsilon_{mouse}$ | **$Day_{ICH}$:[a] -0.24** | **[-0.31, -0.17]** | **1.950E-07** |
| $V_{hema} \sim Day_{ICH}^2 + Day_{ICH} + age + \varepsilon_{mouse}$ | **$Day_{ICH}^2$:[b] 0.02** | **[ 0.01, 0.03]** | **0.002** |
| *Section B: Apparent diffusion coefficient (ADC)* | | | |
| Day 0 and Day 1 data only | | | |
| $ADC_{con}^{MB} \sim Day_{ICH} + age + \varepsilon_{mouse}$ | $Day_{ICH}$:[c] -33.86 | [-67.63, -0.10] | 0.049 |
| $ADC_{con}^{ISO} \sim Day_{ICH} + age + \varepsilon_{mouse}$ | **$Day_{ICH}$: -40.88** | **[-62.50, -19.26]** | **0.002** |
| $ADC_{con}^{HPC} \sim Day_{ICH} + age + \varepsilon_{mouse}$ | $Day_{ICH}$: -28.58 | [-55.47, -1.69] | 0.039 |
| $ADC_{con}^{TH} \sim Day_{ICH} + age + \varepsilon_{mouse}$ | $Day_{ICH}$: -33.73 | [-58.85, -8.61] | 0.013 |
| $ADC_{con}^{HY} \sim Day_{ICH} + age + \varepsilon_{mouse}$ | $Day_{ICH}$: -8.17 | [-58.24, 41.90] | 0.726 |
| $ADC_{con}^{STR} \sim Day_{ICH} + age + \varepsilon_{mouse}$ | $Day_{ICH}$: -18.38 | [-47.41, 10.64] | 0.191 |
| $ADC_{ips}^{MB} \sim Day_{ICH} + age + \varepsilon_{mouse}$ | $Day_{ICH}$: -23.78 | [-68.45, 20.89] | 0.266 |
| $ADC_{ips}^{ISO} \sim Day_{ICH} + age + \varepsilon_{mouse}$ | $Day_{ICH}$: -16.35 | [-43.57, 10.88] | 0.213 |
| $ADC_{ips}^{HPC} \sim Day_{ICH} + age + \varepsilon_{mouse}$ | $Day_{ICH}$: -28.1 | [-51.82, -4.37] | 0.024 |
| $ADC_{ips}^{TH} \sim Day_{ICH} + age + \varepsilon_{mouse}$ | **$Day_{ICH}$: -5.41** | **[-8.60, -2.22]** | **0.003** |
| $ADC_{ips}^{HY} \sim Day_{ICH} + age + \varepsilon_{mouse}$ | $Day_{ICH}$: 2.14 | [-43.96, 48.25] | 0.920 |
| $ADC_{ips}^{STR} \sim Day_{ICH} + age + \varepsilon_{mouse}$ | **$Day_{ICH}$: -183.41** | **[-246.07, -120.75]** | **4.792E-05** |
| $ADC^{MB} \sim Side{\times}Day_{ICH} + age + \varepsilon_{mouse}$ | $Side{\times}Day_{ICH}$:[d] 9.83 | [-30.75, 50.41] | 0.621 |
| $ADC^{ISO} \sim Side{\times}Day_{ICH} + age + \varepsilon_{mouse}$ | $Side{\times}Day_{ICH}$: 15.33 | [-19.30, 49.97] | 0.369 |
| $ADC^{HPC} \sim Side{\times}Day_{ICH} + age + \varepsilon_{mouse}$ | $Side{\times}Day_{ICH}$: -1.42 | [-26.38, 23.55] | 0.908 |
| $ADC^{TH} \sim Side{\times}Day_{ICH} + age + \varepsilon_{mouse}$ | $Side{\times}Day_{ICH}$: 25.5 | [3.79, 47.21] | 0.023 |
| $ADC^{HY} \sim Side{\times}Day_{ICH} + age + \varepsilon_{mouse}$ | $Side{\times}Day_{ICH}$: 9.17 | [-41.43, 59.77] | 0.711 |

| | | | |
|---|---|---|---|
| $ADC^{STR} \sim Side \times Day_{ICH} + age + \varepsilon_{mouse}$ | **Side×$Day_{ICH}$: -177.75** | **[-232.91, -122.59]** | **8.440E-07** |

Day 1, 3, 7, 14, and 21 data only

| | | | |
|---|---|---|---|
| $ADC_{con}^{MB} \sim Day_{ICH} + age + \varepsilon_{mouse}$ | $Day_{ICH}$: 1.34 | [-1.14, 3.82] | 0.277 |
| $ADC_{con}^{ISO} \sim Day_{ICH} + age + \varepsilon_{mouse}$ | $Day_{ICH}$: 0.13 | [-1.70, 1.96] | 0.885 |
| $ADC_{con}^{HPC} \sim Day_{ICH} + age + \varepsilon_{mouse}$ | $Day_{ICH}$: 0.32 | [-1.70, 2.34] | 0.749 |
| $ADC_{con}^{TH} \sim Day_{ICH} + age + \varepsilon_{mouse}$ | $Day_{ICH}$: 0.29 | [-1.73, 2.31] | 0.770 |
| $ADC_{con}^{HY} \sim Day_{ICH} + age + \varepsilon_{mouse}$ | $Day_{ICH}$: 0.66 | [-2.30, 3.61] | 0.652 |
| $ADC_{con}^{STR} \sim Day_{ICH} + age + \varepsilon_{mouse}$ | $Day_{ICH}$: -1.04 | [-2.51, 0.44] | 0.160 |
| $ADC_{ips}^{MB} \sim Day_{ICH} + age + \varepsilon_{mouse}$ | $Day_{ICH}$: 0.98 | [-1.52, 3.48] | 0.430 |
| $ADC_{ips}^{ISO} \sim Day_{ICH} + age + \varepsilon_{mouse}$ | $Day_{ICH}$: -0.36 | [-2.01, 1.29] | 0.658 |
| $ADC_{ips}^{HPC} \sim Day_{ICH} + age + \varepsilon_{mouse}$ | $Day_{ICH}$: 0.53 | [-1.12, 2.20] | 0.516 |
| $ADC_{ips}^{TH} \sim Day_{ICH} + age + \varepsilon_{mouse}$ | $Day_{ICH}$: -0.18 | [-2.40, 2.04] | 0.869 |
| $ADC_{ips}^{HY} \sim Day_{ICH} + age + \varepsilon_{mouse}$ | $Day_{ICH}$: 0.74 | [-1.70, 3.18] | 0.538 |
| $ADC_{ips}^{STR} \sim Day_{ICH} + age + \varepsilon_{mouse}$ | **$Day_{ICH}$: 7.14** | **[4.17, 10.11]** | **3.384E-05** |
| $ADC^{MB} \sim Side \times Day_{ICH} + age + \varepsilon_{mouse}$ | Side×$Day_{ICH}$: -0.44 | [ -2.95, 2.06] | 0.722 |
| $ADC^{ISO} \sim Side \times Day_{ICH} + age + \varepsilon_{mouse}$ | Side×$Day_{ICH}$: -0.54 | [-2.53, 1.44] | 0.587 |
| $ADC^{HPC} \sim Side \times Day_{ICH} + age + \varepsilon_{mouse}$ | Side×$Day_{ICH}$: 0.13 | [ -1.79, 2.04] | 0.896 |
| $ADC^{TH} \sim Side \times Day_{ICH} + age + \varepsilon_{mouse}$ | Side×$Day_{ICH}$: -0.50 | [-2.76, 1.77] | 0.664 |
| $ADC^{HY} \sim Side \times Day_{ICH} + age + \varepsilon_{mouse}$ | Side×$Day_{ICH}$: -0.06 | [ -3.07, 2.95] | 0.969 |
| $ADC^{STR} \sim Side \times Day_{ICH} + age + \varepsilon_{mouse}$ | **Side×$Day_{ICH}$: 9.01** | **[ 6.03, 12.00]** | **1.198E-07** |

*Section C: Cerebral blood perfusion (CBP)*

Day 0 and Day 1 data only

| | | | |
|---|---|---|---|
| $CBP_{con}^{MB} \sim Day_{ICH} + age + \varepsilon_{mouse}$ | **$Day_{ICH}$:[e] -57.93** | **[ -85.95, -29.92]** | **9.690E-04** |
| $CBP_{con}^{ISO} \sim Day_{ICH} + age + \varepsilon_{mouse}$ | **$Day_{ICH}$: -38.95** | **[-55.03, -22.86]** | **3.035E-04** |
| $CBP_{con}^{HPC} \sim Day_{ICH} + age + \varepsilon_{mouse}$ | **$Day_{ICH}$: -36.31** | **[ -49.27, -23.36]** | **9.573E-05** |
| $CBP_{con}^{TH} \sim Day_{ICH} + age + \varepsilon_{mouse}$ | **$Day_{ICH}$: -52.56** | **[-79.18, -25.93]** | **0.001** |
| $CBP_{con}^{HY} \sim Day_{ICH} + age + \varepsilon_{mouse}$ | **$Day_{ICH}$: -50.38** | **[ -65.82, -34.95]** | **2.687E-05** |
| $CBP_{con}^{STR} \sim Day_{ICH} + age + \varepsilon_{mouse}$ | **$Day_{ICH}$: -30.82** | **[-47.91, -13.74]** | **0.002** |
| $CBP_{ips}^{MB} \sim Day_{ICH} + age + \varepsilon_{mouse}$ | **$Day_{ICH}$: -61.91** | **[ -85.76, -38.07]** | **1.766E-04** |
| $CBP_{ips}^{ISO} \sim Day_{ICH} + age + \varepsilon_{mouse}$ | **$Day_{ICH}$: -54.33** | **[ -76.03, -32.63]** | **2.344E-04** |
| $CBP_{ips}^{HPC} \sim Day_{ICH} + age + \varepsilon_{mouse}$ | **$Day_{ICH}$: -44.00** | **[ -57.76, -30.25]** | **3.192E-05** |
| $CBP_{ips}^{TH} \sim Day_{ICH} + age + \varepsilon_{mouse}$ | **$Day_{ICH}$: -46.97** | **[ -78.47, -15.47]** | **7.720E-03** |
| $CBP_{ips}^{HY} \sim Day_{ICH} + age + \varepsilon_{mouse}$ | **$Day_{ICH}$: -55.24** | **[ -72.12, -38.37]** | **2.619E-05** |
| $CBP_{ips}^{STR} \sim Day_{ICH} + age + \varepsilon_{mouse}$ | **$Day_{ICH}$: -66.92** | **[ -77.53, -56.32]** | **6.482E-08** |

| | | | |
|---|---|---|---|
| $CBP^{MB} \sim Side{\times}Day_{ICH} + age + \varepsilon_{mouse}$ | $Side{\times}Day_{ICH}$:[f] -3.95 | [ -32.25, 24.34] | 0.774 |
| $CBP^{ISO} \sim Side{\times}Day_{ICH} + age + \varepsilon_{mouse}$ | $Side{\times}Day_{ICH}$: -16.51 | [ -35.72, 2.70] | 0.088 |
| $CBP^{HPC} \sim Side{\times}Day_{ICH} + age + \varepsilon_{mouse}$ | $Side{\times}Day_{ICH}$: -7.47 | [-21.47, 6.53] | 0.280 |
| $CBP^{TH} \sim Side{\times}Day_{ICH} + age + \varepsilon_{mouse}$ | $Side{\times}Day_{ICH}$: 6.82 | [-26.23, 39.88] | 0.672 |
| $CBP^{HY} \sim Side{\times}Day_{ICH} + age + \varepsilon_{mouse}$ | $Side{\times}Day_{ICH}$: -4.34 | [ -22.39, 13.71] | 0.622 |
| $CBP^{STR} \sim Side{\times}Day_{ICH} + age + \varepsilon_{mouse}$ | **$Side{\times}Day_{ICH}$: -35.04** | **[ -52.05, -18.04]** | **3.290E-04** |
| Day 1, 3, 7, 14, and 21 data only | | | |
| $CBP_{con}^{MB} \sim Day_{ICH} + age + \varepsilon_{mouse}$ | $Day_{ICH}$: 0.74 | [ -0.59, 2.07] | 0.262 |
| $CBP_{con}^{ISO} \sim Day_{ICH} + age + \varepsilon_{mouse}$ | $Day_{ICH}$: 0.60 | [ -0.97, 2.16] | 0.440 |
| $CBP_{con}^{HPC} \sim Day_{ICH} + age + \varepsilon_{mouse}$ | $Day_{ICH}$: 0.63 | [-0.53, 1.79] | 0.276 |
| $CBP_{con}^{TH} \sim Day_{ICH} + age + \varepsilon_{mouse}$ | $Day_{ICH}$: 0.71 | [-0.66, 2.08] | 0.294 |
| $CBP_{con}^{HY} \sim Day_{ICH} + age + \varepsilon_{mouse}$ | $Day_{ICH}$: 0.72 | [-0.55, 1.99] | 0.257 |
| $CBP_{con}^{STR} \sim Day_{ICH} + age + \varepsilon_{mouse}$ | $Day_{ICH}$: 0.46 | [-0.56, 1.48] | 0.366 |
| $CBP_{ips}^{MB} \sim Day_{ICH} + age + \varepsilon_{mouse}$ | $Day_{ICH}$: 0.95 | [ -0.38, 2.28] | 0.152 |
| $CBP_{ips}^{ISO} \sim Day_{ICH} + age + \varepsilon_{mouse}$ | $Day_{ICH}$: 0.77 | [ -0.88, 2.43] | 0.346 |
| $CBP_{ips}^{HPC} \sim Day_{ICH} + age + \varepsilon_{mouse}$ | $Day_{ICH}$: 0.74 | [ -0.46, 1.94] | 0.214 |
| $CBP_{ips}^{TH} \sim Day_{ICH} + age + \varepsilon_{mouse}$ | $Day_{ICH}$: 0.56 | [ -1.35, 2.47] | 0.551 |
| $CBP_{ips}^{HY} \sim Day_{ICH} + age + \varepsilon_{mouse}$ | $Day_{ICH}$: 0.87 | [-0.60, 2.33] | 0.236 |
| $CBP_{ips}^{STR} \sim Day_{ICH} + age + \varepsilon_{mouse}$ | **$Day_{ICH}$: 3.75** | **[ 2.81, 4.69]** | **1.042E-08** |
| $CBP^{MB} \sim Side{\times}Day_{ICH} + age + \varepsilon_{mouse}$ | $Side{\times}Day_{ICH}$: 0.02 | [ -1.47, 1.52] | 0.975 |
| $CBP^{ISO} \sim Side{\times}Day_{ICH} + age + \varepsilon_{mouse}$ | $Side{\times}Day_{ICH}$: 0.19 | [ -1.71, 2.09] | 0.843 |
| $CBP^{HPC} \sim Side{\times}Day_{ICH} + age + \varepsilon_{mouse}$ | $Side{\times}Day_{ICH}$: 0.06 | [ -1.30, 1.42] | 0.930 |
| $CBP^{TH} \sim Side{\times}Day_{ICH} + age + \varepsilon_{mouse}$ | $Side{\times}Day_{ICH}$: 0.02 | [ -1.95, 2.00] | 0.980 |
| $CBP^{HY} \sim Side{\times}Day_{ICH} + age + \varepsilon_{mouse}$ | $Side{\times}Day_{ICH}$: 0.29 | [-1.32, 1.90] | 0.719 |
| $CBP^{STR} \sim Side{\times}Day_{ICH} + age + \varepsilon_{mouse}$ | **$Side{\times}Day_{ICH}$: 3.31** | **[ 2.14, 4.47]** | **5.527E-07** |

[a]Unit of $Day_{ICH}$ effect in hematoma volume: μL/day;

[b]Unit of $Day_{ICH}^2$ effect in ADC: $\mu L/day^2$;

[c]Unit of $Day_{ICH}$ effect in ADC: $\times 10^{-6}$ $mm^2/s/day$;

[d]Unit of $Side{\times}Day_{ICH}$ effect in ADC: $\times 10^{-6}$ $mm^2/s/day$;

[e]Unit of $Day_{ICH}$ effect in CBP: ml/100g/min/day;

[f]Unit of $Side{\times}Day_{ICH}$ effect in CBP: ml/100g/min/day.

[g]*p*-values were reported in scientific notation when appropriate. For example, 1.950E-07 represents $1.950 \times 10^{-7}$.
